# Evaluating the impact of NPC1 SNPs on entry efficiency of Filovirus *in vitro*: agent-based model approach

**DOI:** 10.1101/2025.03.13.642967

**Authors:** Ju Seong Kim, Kwang Su Kim, Ayato Takada, Yusuke Asai, Shingo Iwami, Seung-Woo Son, Mi Jin Lee

## Abstract

The interaction between the Niemann-Pick C1 (NPC1) protein and the glycoprotein (GP) of filoviruses is essential for viral entry into host cells. Single nucleotide polymorphisms (SNPs) in NPC1, which lead to amino acid substitutions, can significantly alter viral entry efficiency. However, the quantitative impact of these SNPs remains unclear. To address this, we investigate in vitro cell-to-cell infection using a plaque assay with vesicular stomatitis virus (VSV) expressing Ebola and Marburg GP. We employe an agent-based model (ABM) to estimate entry efficiency by simulating plaque growth, enabling a comparative analysis of SNP-specific effects on viral entry. Our quantitative analysis reveals that D508N, P424A, and S425L substitutions in NPC1 reduce entry efficiency in both viruses. These findings provide insights into host-pathogen interactions and demonstrate the value of ABM in virology research, potentially guiding therapeutic strategies targeting viral entry mechanisms.

**Author summary:** Ebola and Marburg viruses are highly pathogenic filoviruses that cause severe hemorrhagic fever in humans, with case fatality rates reaching around 50%. These viruses pose significant public health challenges due to their potential for large-scale outbreaks. A key step in their infection process is the interaction between the Niemann-Pick C1 (NPC1) protein on host cells and the viral glycoprotein, which facilitates viral entry. Genetic variations in NPC1 caused by single nucleotide polymorphisms (SNPs) can lead to amino acid substitutions, potentially altering the efficiency of viral entry. To better understand this process, we developed an agent-based model, a computational approach that simulates individual cell interactions and provides spatial resolution not achievable with traditional modeling techniques, to simulate viral plaque growth *in vitro*. By applying this model, we quantified how specific NPC1 substitutions, such as D508N, P424A, and S425L, affect the entry efficiency of Ebola and Marburg viruses. Our findings highlight the potential of computational modeling to uncover the impact of genetic variations on viral infections and provide insights that may inform therapeutic strategies against these deadly viruses.

## Introduction

Filoviruses, including Ebolavirus (EBOV) and Marburgvirus (MARV), cause severe hemorrhagic fever in humans, with an average case fatality rate of approximately 50%. Despite significant advances in vaccine development in recent years, currently approved vaccines are primarily limited to the Zaire strain of Ebola. Vaccines targeting other EBOV strains and MARV remain at various stages of development, with several candidates undergoing clinical trials but none yet widely approved for general use [1]. These viruses have been responsible for large-scale outbreaks, predominantly in Africa, and their potential spread to other regions poses a serious global public health threat [2]. As a result, the World Health Organization (WHO) has designated filovirus research and response efforts as a critical priority [3]. Extensive studies have been conducted to elucidate the infection mechanisms of these viruses, with a particular focus on key stages of viral entry, replication, and interactions with host cellular components [4–7].

Filoviruses enter host cells through receptor-mediated endocytosis, where glycoproteins (GPs) on the viral envelope bind to receptors on the host cell surface such as TIM-1 or C-type lectins. Upon entry, the viruses are transported to late endosomes, where GPs undergo sequential cleavage by cathepsins B and L, leading to the formation of digested GP (dGP). The acidic environment of the late endosome triggers membrane fusion between dGP and Niemann-Pick C1 (NPC1), facilitating the release of viral RNA into the cytoplasm, where replication and dissemination occur [8– 10]. Recent studies have provided deeper insights into the structural basis of this interaction, with cryo-electron microscopy revealing GP conformational changes that enhance NPC1 binding under endosomal conditions. These findings have refined our understanding of the molecular mechanisms governing filovirus entry, expanding upon earlier models of the process.

Previous studies have highlighted the critical role of NPC1 in mediating viral entry. Côté et al. demonstrated that small-molecule inhibitors targeting NPC1 effectively block its interaction with the virus, hereby preventing Ebola virus infection [11]. Furthermore, a genome-wide haploid genetic screen conducted by Carette et al. showed the importance of NPC1 by identifying its interaction with the viral GP [12]. Single-nucleotide polymorphisms (SNPs) represent genetic variations that can alter amino acid sequences, potentially altering the structure and function of critical proteins involved in viral entry pathways. These structural alterations may directly affect the binding interface between NPC1 and viral GPs, influencing binding affinity, kinetics, or the stability of the protein-protein interaction. Kondoh et al. demonstrated that SNP-induced amino acid substitutions on NPC1 significantly affect entry efficiency by examining a candidate list of naturally occurring SNPs [13]. However, the quantitative effects of SNP-induced changes in entry efficiency remain poorly understood. Viral entry is a complex, multifactorial process, making it challenging to experimentally control all variables to determine the effects of SNPs.

Therefore, modeling provides a valuable approach to quantitatively comparing the impact of different SNPs. Previous studies have attempted to analyze plaque (the cluster consisting of dead cells) assay quantitatively using an ordinary differential equation (ODE) approach [14-17]. However, Conventional ODE models assume a well-mixed system, where infection occurs independently of spatial constraints. This assumption oversimplifies real-world infection dynamics and fails to account for phenomena such as saturation effects. Infection rates and dynamics are often influenced by the spatial distribution of host cells. To overcome these limitations, several efforts have been made to incorporate spatial information into ODE models [18-20]. Despite these advancements, ODE models still struggle to fully capture the complexity of real-world infection dynamics, particularly at the level of individual cell interactions where stochastic processes and spatial arrangements significantly influence viral spread.

In this paper, we focus on comparing the viral entry efficiency of vesicular stomatitis virus (VSV) ΔG-EBOV and VSV ΔG-MARV as they interact with Vero E6 cells expressing human NPC1 during cell-to-cell infection. To quantitatively assess the effects of different SNPs in NPC1 on entry efficiency, we developed an agent-based model (ABM) of plaque growth. While previous studies, such as [18], have attempted similar analyses using ordinary differential equation (ODE) approaches, these models are limited by their assumption of a well-mixed state and lack of spatial resolution. In contrast, our ABM approach not only incorporates spatial dynamics but also introduces stochasticity into the modeling process [21-24]. The stochastic nature of ABM offers distinct advantages over traditional modeling approaches. By incorporating probabilistic interactions at the individual agent level, our ABM captures the complexity and variability inherent in real-world infection dynamics. This stochasticity enables more realistic simulations of viral spread and cell-to-cell interactions, providing a nuanced understanding of filovirus infection processes. Consequently, our methodology provides a more nuanced understanding of filovirus infection processes. Moreover, our ABM framework is adaptable for studying other viruses with available plaque data, offering a versatile tool for investigating both spatial and stochastic infection dynamics.

## Results

### Agent-based plaque growing model

To indirectly compare the entry efficiency of Ebola and Marburg viruses affected by SNP-induced mutations in NPC1, we utilize an agent-based model (ABM) for the plaque (dead-cell cluster) growth. By simulating plaque growth, we measure the plaque radius and compare it with experimentally obtained values from plaque assays. This approach allows us to estimate entry efficiency based on model-calibrated parameters. Since our focus is on the effects of SNPs on entry efficiency, we analyze only a single plaque radius from plaque assay in **Fig 1A** and **Fig A** in **S1 Text**. In our plaque assay experiments, cells are placed in a monolayer on a plate and an agar-containing medium is used to immobilize cell movement to ensure that viral spread occurs exclusively through cell-to-cell transmission. To accurately reflect this setup, we assume a triangular lattice arrangement for the cells. In **Fig 2A**, the cells are depicted as hexagons, highlighting the six neighbors each cell has within the triangular lattice.

**Fig 1.**
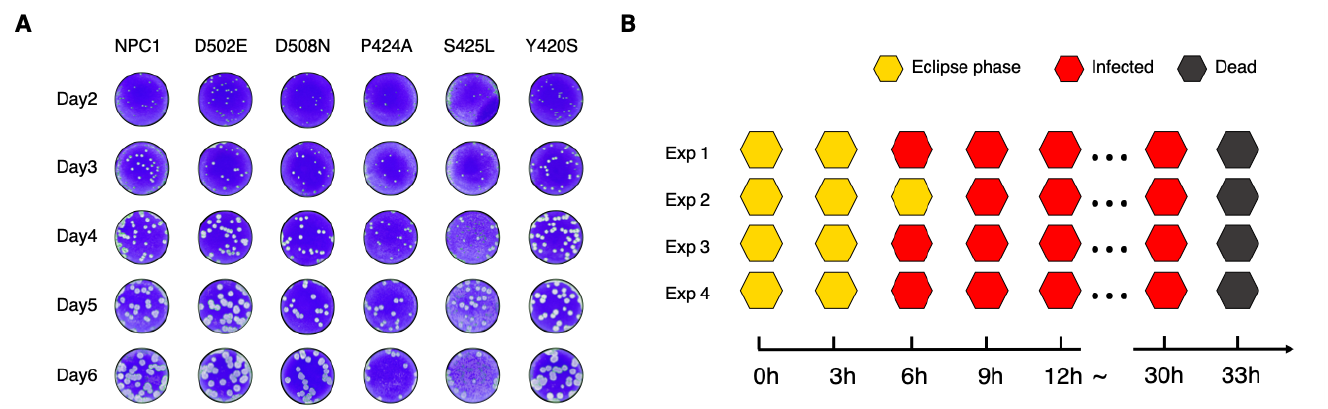
Plaque assay & Virus production assay with VSVΔG-EBOV: (A) Plaque formation of VSVΔG-EBOV on Vero E6 cells expressing wild-type NPC1 and five SNP mutants (D502E, D508N, P424A, S425L, and Y420S) observed from days 2 to 6. The White areas represent plaques formed by dead cells. (B) Virus production assays using Vero E6/NPC1-KO cells expressing human NPC1, observed starting immediately after viral infection. Data were collected in four independent experiments, with cell states recorded every three hours. The recorded states include the eclipse phase cells (yellow), infected cells (red), and dead cells (gray) over a 33-hour period.

**Fig 2.**
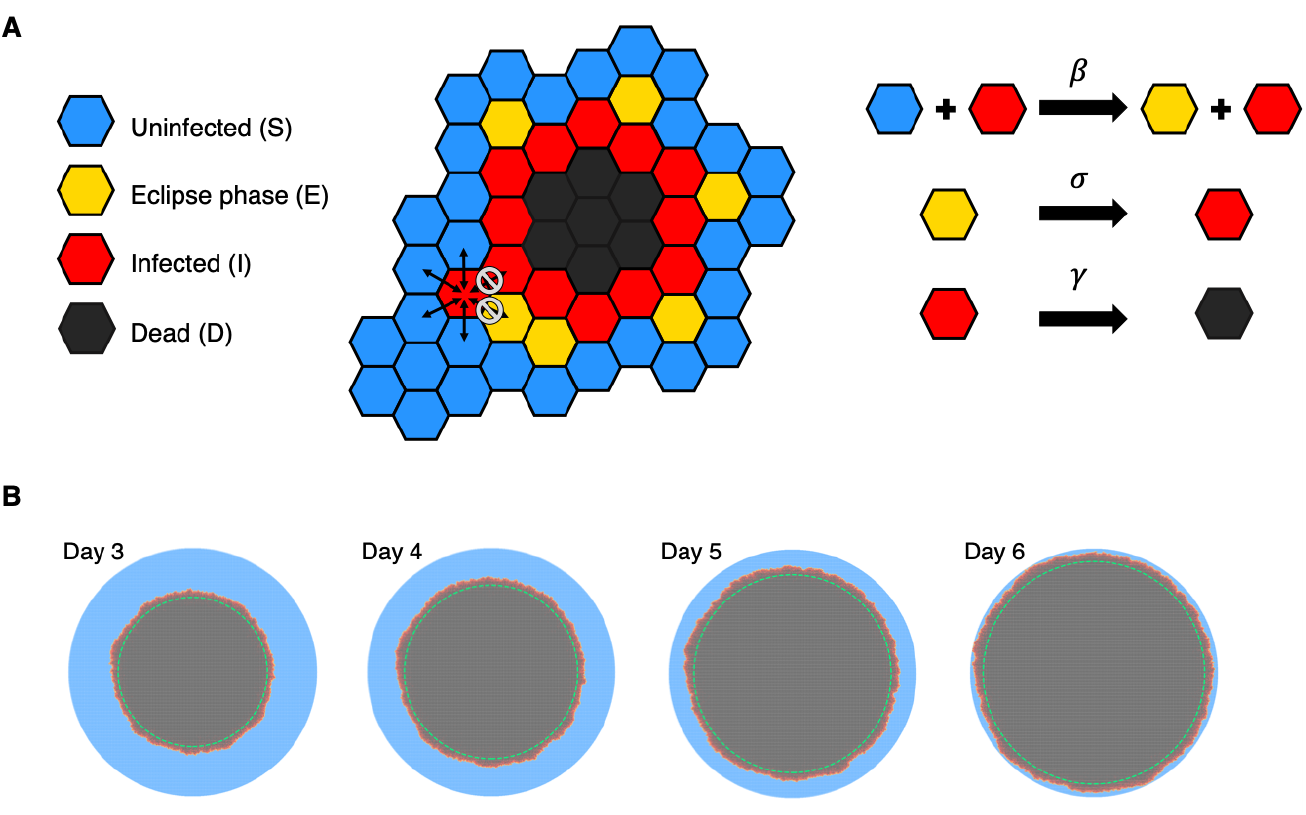
Modeling and plaque visualization of cell configurations: (A) Schematic representation of viral spread dynamics within a triangular lattice of cells. Infected cells (red) can transmit the virus to neighboring uninfected cells (blue) with probability *β* and transition to dead state (black) with probability *γ*. Cells in the eclipse phase (yellow) do not interact with other cells but become infectious with probability *σ*. (B) Simulated plaque growth from day 3 to day 6 in one-day intervals. The dashed green circles represent the estimated plaque size.

In our model, each cell is classified into one of four distinct states: uninfected or susceptible (S), eclipse phase (E), infected (I), or dead (D). Cells transition sequentially through these states, starting from uninfected to eclipse phase, then to infected, and finally to dead, as

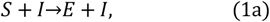

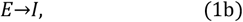

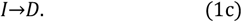

Infected cells can transmit the virus to neighboring uninfected cells, as described in **Eq (1a)**. This model requires three key parameters to govern state transitions: *β, σ*, and *γ*. Here, *β* represents the transmission rate of the infectious virus, *σ* represents the transition rate from the eclipse phase (E) to the infected (infectious) phase (I), and *γ* denotes the rate from the infected phase (I) to dead (D). We assume SNPs do not affect the duration of the eclipse or infected phases. The values for *σ* and *γ* are inferred from the durations of the eclipse and infected phases, respectively, based on data from the virus production assay shown in **Fig 1B**. Given that three of the four independent experiments yielded eclipse phase durations of 3 to 6 hours and the infectious durations of 24 to 30 hours, we selected the median values within these ranges, using *σ* = 1/4.5h and *γ* = 1/27h as representative estimates. Furthermore, to accurately estimate *β*, the plaques are grown from a single seed until it reaches the radius observed on the day 2 of the plaque assay for each SNP, which is then used as the initial plaque radius.

### Plaque radius estimation in model

The estimated plaque radius 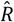 is determined from the average radius of a single cell (*r*) and the number of dead cells (*n*). By equating the area of a large circle to the sum of the areas of smaller circles, the radius of the large circle is approximated as

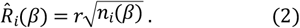

The estimation is based on the assumption that the dead cells expand radially in a fully packed assay under cell-to-cell infection, as the infected cells select their uninfected neighbors in an isotropic manner. From day 2 to day 6, the green dashed circle in NPC1 non-mutated cases represents the estimated plaque boundary (**Fig 2B**). Although this method slightly underestimates the actual plaque size, its effect on the comparative analysis of SNPs is negligible. Since the number of plaque radius data points on day *j* (*n*_*j*_) for each SNP varies and is relatively small, we use all available data points to calculate the error values in

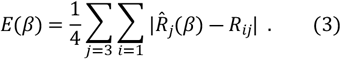

Here, *R*_*ij*_ represents the radius of the *i*-th plaque measured on day *j* from empirical data. This allows us to account for the variability in plaque measurements across different days and SNPs. **Fig B and Fig C** in **S1 Text** represent the error function as a function of the variable. Thus, we define heuristically the range of *β* values as those falling within 20% of the minimum error value, 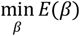. For each SNP, the optimal *β* value is determined using the error function described above, and the corresponding optimal *β* values are presented in **Fig D** in **S1 Text**.

### Entry efficiency comparison

To evaluate the impact of SNP-induced mutations on viral entry efficiency, we used the wild-type NPC1 protein as a baseline. The relative entry efficiency for each SNP is defined as *α* = *β*_SNP_/*β*_NPC1_, where *β*_SNP_ represents the transmission rate associated with a specific SNP. For both Ebola and Marburg viruses, the D052E and Y420S mutations exhibited *α* values greater than 1, indicating enhanced entry efficiency compared to wild-type NPC1. In contrast, D508N, P424A and S425L SNPs showed *α* values less than 1, suggesting reduced entry efficiency. Notably, the P424A mutation in the Ebola virus resulted in the most significant reduction, with an *α* value decreasing by 47%, making it the SNP with the greatest inhibitory effect on entry. In addition, Marburg virus, consistently displayed higher *α* value than Ebola virus, implying that SNP-induced variations in NPC1 may have a more pronounced effect on Marburg virus entry (**Fig 3A** and **Fig 3B**).

**Fig 3.**
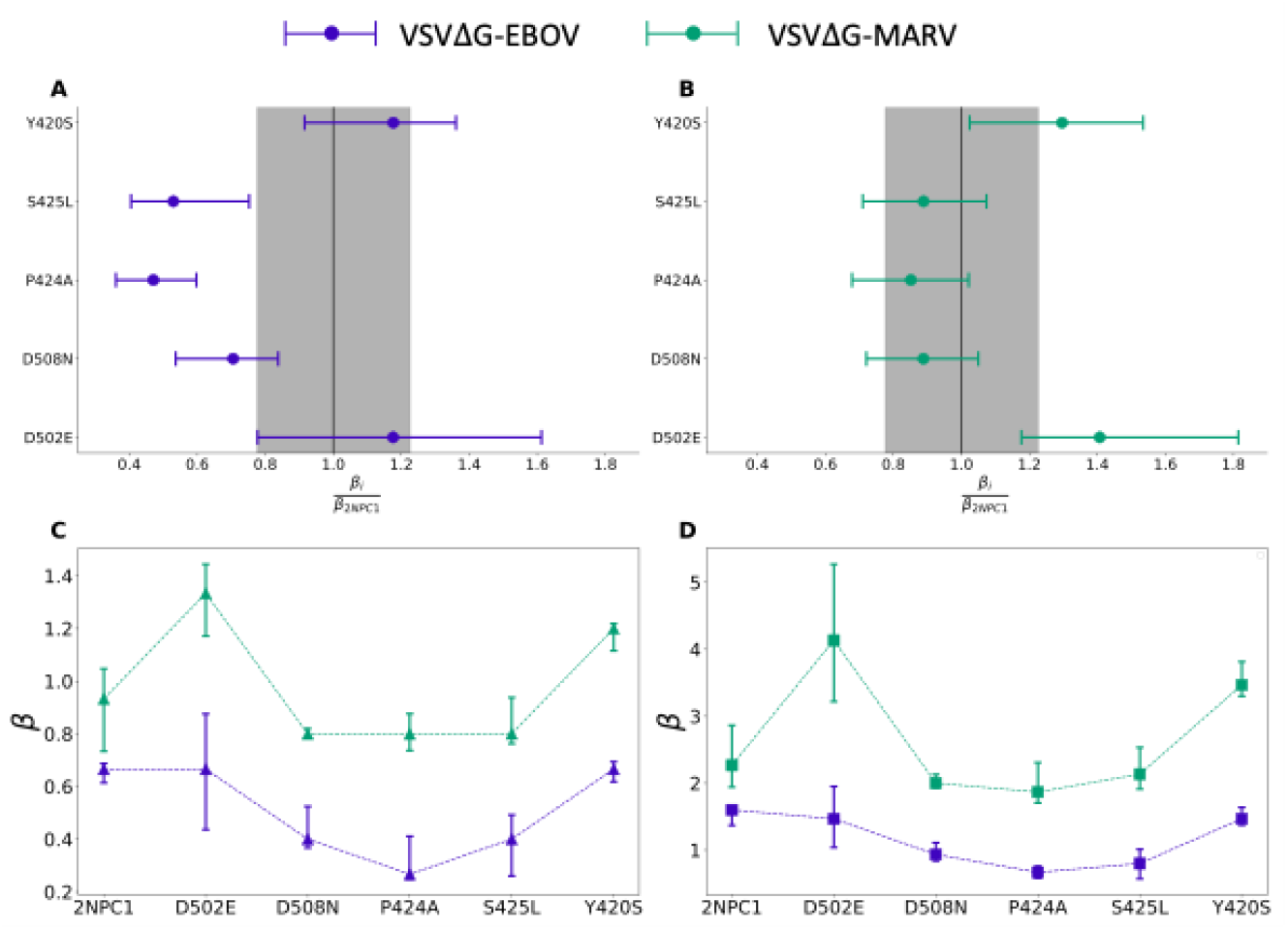
Entry efficiency comparison: Comparison of entry efficiency between VSVΔG-EBOV (A) and VSVΔG-MARV (B). Entry efficiency is calculated as the ratio of the efficiency for each SNP to the wild-type NPC1. The central point represents the value obtained using the *β* parameter that minimize the error, while the error bars indicate the range of possible *β* within 20% of the minimum error. The gray-shaded area represents the range corresponding to wild-type NPC1, serving as a reference for comparison. The panels (C) and (D) show the optimal *β* values for Ebola virus and Marburg virus, with square markers representing the square lattice and triangular markers representing the triangular lattice.

### Lattice structure dependency

To simulate the infection process in the fully packed assay, we initially assume a triangular lattice. To evaluate how *β* varies with different lattice structures, we have conducted simulations on a square lattice for both viruses. The lattice structure determines the number of neighboring cells each cell interacts with. The triangular lattice has six neighboring cells, whereas the square lattice has four neighboring cells. To achieve plaque sizes consistent with experimental observations, a higher infection rate is necessary for the square lattice compared to the triangular lattice. This is because, with fewer neighboring cells in the square lattice (4 versus 6), each cell has fewer opportunities for viral transmission, requiring a higher per-contact transmission rate to achieve the same overall spread. Consequently, **Fig 3C and Fig 3D** show that *β* is consistently higher for the square lattice in all regions. Although the exact number of neighbors that each biological cells interacts with remains uncertain, our results indicate that while the absolute *β* value varies with the number of neighboring cells, the relative ranking of *β* values among different SNPs remains largely unchanged.

## Discussion

Using the ABM approach, we observed that the entry efficiency for VSVΔG-EBOV ranged from -47% to +18%, whereas for VSVΔG-MARV, it varied from -15% to +41%. These results suggest that NPC1 substitutions generally lead to a significant reduction in entry efficiency for VSVΔG-EBOV, with relatively smaller increases, whereas the opposite trend was observed for VSVΔG-MARV. These quantitative differences highlight the virus-specific effects of NPC1 mutations, which could inform the development of targeted antiviral strategies. Notably, substitutions such as D508N, P424A, and S425L reduced the entry efficiency in both viruses, whereas D508N and Y420S increased the efficiency in both cases. These findings align with previous research on EBOV [13], while revealing subtle variations for MARV, underscoring the ability of ABM to capture finer differences in infection dynamics compared to ordinary differential equation (ODE) models.

Mathematical models used to analyze plaque assays encompass various approaches, including the diffusion model [25-27] and the ODE model [19, 20]. Although these models provide valuable frameworks for understanding viral spread, they rely on continuous equations and population-level dynamics, making them less suited for capturing discrete cell-to-cell interactions. In contrast, ABM offers greater flexibility by explicitly representing individual agents, their interactions, and their environments. In some cases, new cells are continuously generated. Although the present model does not account for cell regeneration, a more realistic simulation could be achieved by extending the constructed model to include a mechanism whereby dead cells are replaced by susceptible cells [30]. The spatial arrangement of cells can significantly influence the outcomes of plaque assays [20, 28]. Our model assumes a lattice structure for the fully packed cell placement in the assay; however, incorporating off-lattice configurations could offer a more biologically realistic representation of cellular interaction [24]. Several studies have demonstrated that ABM can be extended into three-dimensional environments to capture more realistic spatial and structural dynamics [30, 33]. In addition, some advanced models have incorporated factors such as gravitational effects on cell positioning, introducing an additional layer of complexity to the simulation [32].

While our ABM provides valuable insights into the effects of NPC1 mutations on filovirus entry efficiency, it has several limitations. The model simplifies the complexity of the viral entry process by focusing solely on GP-to-NPC1 interactions, excluding other host factors and intracellular processes that may influence viral replication and spread. In addition, the analysis is constrained by the use of daily data intervals and the selection of only non-merged plaques. While necessary for measurement consistency, these constraints may reduce dataset representativeness and limit the resolution of entry efficiency comparisons. Increasing the frequency of data collection or expanding the dataset with multiple independent experiments could enhance the measurement precision and enable more robust statistical analysis.

The high computational complexity of ABM imposes limitations on extending simulations to longer time scales or more diverse environmental conditions, as assigning individual states to each cell and performing detailed calculations require substantial computational resources. Despite these constraints, ABM-based approach provides valuable insights into individual susceptibility and infection dynamics. By explicitly modeling cell-to-cell interactions, it serves as a powerful framework for studying viral entry mechanisms and infection spread.

The adaptability of ABM approach enables its application to plaque assays across various viral infections, providing a valuable tool for studying host-pathogen interactions beyond filoviruses. Furthermore, this method can aid in prioritizing treatment for high-risk individuals and optimizing the allocation of limited resources, such as vaccines, thereby supporting more informed and effective decision-making in public health and clinical settings.

## Methods

### Plaque assay & Virus production assay

Vero E6 cells, which are African green monkey kidney epithelial cells, were infected with VSVΔG-EBOV, as shown in **Fig 1A**. The cells were seeded as a monolayer and immobilized using an agar-containing minimum essential medium Eagle. This ensure that the infection rate measurement specifically reflected cell-to-cell transmission. To maintain a consistent initial cell population, the medium was adjusted to prevent cell proliferation. For analysis, only non-merged plaques are selected to ensure accurate measurements of the plaque radius. In a separate assay, four independent experiments were conducted for both VSVΔG-EBOV and VSVΔG-MARV to investigate the duration of virus status, with measurements taken at three-hour intervals (**Fig 1B)**. The cellular infection was assessed based on cytopathic effects. The data from these assays are identical to those used in the previous paper, where a more detailed description can be found in the previous paper [17].

## LIST OF SUPPLEMENTARY MATERIALS

**S1 Text**. Fig A. VSV G-MARV plaque assay and virus production assay. Fig B: Error function for VSVΔ-EBOV infected plaque with NPC1 error function is shown. The blue shaded area represents the range of *β* values within 20% of the minimum error value. Fig C. Error function for VSVΔ-MARV infected plaque with NPC1 error function is shown. The blue shaded area represents the range of *β* values within 20% of the minimum error value. Fig D. Best fitting of *β* each SNPs. The simulation-derived plot shows the radius of the plaque against the optimal *β*. Video A. Dynamics of the single seed infection process. Video B. Dynamics of multi-seed infection process.

This work was supported by the National Research Foundation of Korea (NRF) grant funded by the Korea government (MSIT) through Grant Numbers. 2022R1C1C2003637 (K.S.K.), NRF-2023R1A2C1007523 (S.-W.S.), and RS-2024-00341317 (M.J.L.), MIRAI JPMJMI22G1 (S.I.), Moonshot R&D JPMJMS2021 (S.I.) and JPMJMS2025 (S.I.), This work was also partly supported by the Institute of Information & communications Technology Planning & Evaluation (IITP) grant funded by the Korean government (MSIT) (No.RS-2022-00155885, Artificial Intelligence Convergence Innovation Human Resources Development (Hanyang University ERICA)).

